# The Rumen Metatranscriptome Landscape Reflects Dietary Adaptation and Methanogenesis in Lactating Dairy Cows

**DOI:** 10.1101/275883

**Authors:** Bastian Hornung, Bartholomeus van den Bogert, Mark Davids, Vitor A.P. Martins dos Santos, Caroline M. Plugge, Peter J. Schaap, Hauke Smidt

## Abstract

Methane eructed by ruminant animals is a main contributor to greenhouse gas emissions and is solely produced by members of the phylum *Euryarchaeota* within the domain *Archaea*. Methanogenesis depends on the availability of hydrogen, carbon dioxide, methanol and acetate produced, which are metabolic products of anaerobic microbial degradation of feed-derived fibers. Changing the feed composition of the ruminants has been proposed as a strategy to mitigate methanogenesis of the rumen microbiota.

We investigated the impact of corn silage enhanced diets on the rumen microbiota of rumen-fistulated dairy cows, with a special focus on carbohydrate breakdown and methanogenesis. Metatranscriptome analysis of rumen samples taken from animals fed corn silage enhanced diets revealed that genes involved in starch metabolism were significantly more expressed while archaeal genes involved in methanogenesis showed lower expression values. The nutritional intervention also influenced the cross-feeding between *Archaea* and *Bacteria*.

The results indicate that the ruminant diet is important in methanogenesis. The diet-induced changes resulted in a reduced methane emission. The metatranscriptomic analysis provided insights into key underlying mechanisms and opens the way for new rational methods to further reduce methane output of ruminant animals.

## Introduction

Reduction of global greenhouse gas (GHG) output is necessary to prevent a further increase in global warming, which is predicted to result in multiple detrimental effects for the environment and human affairs (Schleussner et al., 2016). The necessary measures are focused on the industrial and agricultural sectors in developed countries, with the aim to reduce carbon dioxide, methane and other GHG emissions. One of the predominant sources of methane emission, estimated to be as high as ~35% of the total anthropogenic methane emissions worldwide (IPCC, 2007;McMichael et al., 2007), is the agricultural sector, and especially the eructation by ruminant animals (Murray et al., 2007).

Ruminal microbes play a pivotal role in the breakdown of animal feed and contribute between 35 to 50% of the animal’s energy intake (Bergman, 1990). The ruminal microbial composition is complex, with diverse populations including bacteria, archaea, fungi, and protozoa. Their functional capacity is vast and has not yet been fully elucidated (Hess et al., 2011; Li et al., 2012).

Notwithstanding the ruminal microbial complexity, methane is solely produced by a few members of the phylum *Euryarcheota* belonging to the *Archaea* (Hook et al., 2010). It has been shown that a change in diet can have a significant effect on the methane emissions of ruminants (Liu et al., 2011;van Gastelen et al., 2015), but the mechanisms that drive this change are not fully understood. The methanogenic archaea are not directly involved in the breakdown of the feed, but rely on their relationships with other community members that provide the necessary substrates for methanogenesis like hydrogen, formate and methanol. Microbial ecology in cows and other ruminants has been investigated using 16S ribosomal RNA (rRNA) genes as molecular markers (Fernando et al., 2010;Pitta et al., 2014), the sheep rumen microbial metatranscriptome has been investigated (Shi et al., 2014), and in cows specialized and general microbial functions have been examined (Brulc et al., 2009Hess et al., 2011;Poulsen et al., 2013;Dai et al., 2014;Dassa et al., 2014;Roehe et al., 2016;Jose et al., 2017). Understanding the mechanisms that influence cow rumen methanogenesis requires community-level analysis of active metabolic functions, however, a comprehensive analysis of diet-dependent effects on the functional landscape of the rumen microbiota is lacking. Here we investigated the effect of feed composition on bovine rumen activity patterns with a special focus on methane metabolism. By analysis of the rumen metatranscriptome landscapes in animals fed mixed grass silage (GS) and corn silage (CS) diets, we were able to elucidate the impact of the diet on the expression of methanogenic pathways and on the relationships of methanogens with other community members.

## Materials and Methods

### Study design and sampling

The study design has been described in detail by Van Gastelen et al. (van Gastelen et al., 2015). Briefly, the experiment was performed in a complete randomized block design with four dietary treatments and 32 multiparous lactating Holstein-Friesian cows. Cows were blocked according to lactation stage, parity, milk production, and presence of a rumen fistula (12 cows). Within each block cows were randomly assigned to 1 of 4 dietary treatments. All dietary treatments had a roughage-to-concentrate ratio of 80:20 based on dry matter. In the four diets, the roughage consisted of either 100% GS (**GS100**), 67% GS and 33% CS (**GS67**), 33% GS and 67% CS (**GS33**), or 100% CS (**GS0**; all dry matter basis). This study, including the rumen fluid sampling, was conducted in accordance with Dutch law and approved by the Animal Care and Use Committee of Wageningen University.

### Sample collection and processing

In total, samples from 12 rumen fistulated cows, three per dietary treatment, were used for metatranscriptome analysis. Rumen fluid was collected 3 hours after morning feeding on day 17 of the experimental period (for further details regarding the whole experimental period, see (van Gastelen et al., 2015)). The samples were obtained as described previously (van Zijderveld et al., 2011), and collected from the middle of the ventral sac. The rumen fluid samples were immediately frozen on dry ice and subsequently transported to the laboratory where the samples were stored at -80° C until further analysis.

For RNA extraction, 1 ml rumen fluid was centrifuged for 5 min at 9000 g, after which the pellet was re-suspended in 500 μl TE buffer (Tris-HCl pH 7.6, EDTA, pH 8.0). Total RNA was extracted from the resuspended pellet according to the Macaloid-based RNA isolation protocol (Zoetendal et al., 2006) with the use of Phase Lock Gel heavy (5 Prime GmbH, Hamburg) (Murphy and Hellwig, 1996) during phase separation. The aqueous phase was purified using the RNAeasy mini kit (Qiagen, USA), including an on-column DNAseI (Roche, Germany) treatment as described previously (Zoetendal et al., 2006). Total RNA was eluted in 30 μl TE buffer. RNA quantity and quality were assessed using NanoDrop ND-1000 spectrophotometer (Nanodrop Technologies, Wilmington, USA) and Experion RNA Stdsens (Biorad Laboratories Inc., USA).

rRNA was removed from the total RNA samples using the Ribo-Zero™ rRNA removal Kit (Meta-Bacteria; Epicentre, Madison, WI, USA) using 5 μg total RNA as input. Subsequently, barcoded cDNA libraries were constructed for each of the rRNA depleted samples using the ScriptSeq™ Complete Kit (Bacteria; Epicentre) according to manufacturer’s instructions in combination with Epicentre’s ScriptSeq Index PCR Primers.

The barcoded cDNA libraries were pooled and sent to GATC Biotech (Konstanz, Germany) for 150 bp single end sequencing on one single lane using the Illumina HiSeq2500 platform in combination with the TruSeq Rapid SBS (200 cycles) and TruSeq Rapid SR Cluster Kits (Illumina Inc., San Diego, CA, USA).

### Bioinformatics

The general workflow for data quality assessment and filtering was adapted from (Davids et al., 2016). rRNA reads were removed with SortMeRNA v1.9 (Kopylova et al., 2012) and all included databases. Adapters were trimmed with cutadapt v1.2.1 (Martin, 2011) using default settings except for an increased error value of 20 % for the adapters. The latter was chosen considering that with the default setting of 10% adapter sequences could still be found after trimming. Quality trimming was performed with PRINSEQ Lite v0.20.0 (Schmieder and Edwards, 2011) with a minimum sequence length of 40 bp and a minimum quality of 30 at both ends of the read and as mean quality. All reads with non-IUPAC characters were discarded as were all reads containing more than three Ns. Details on the RNAseq raw data analysis can be found in Supplementary Table 1. The log files with the used commands can be found in supplementary file 1 and the used python script in supplementary file 2. The raw data was deposited at EBI ENA, and can be accessed under accession numbers ERS685245 - ERS685256.

### Assembly and annotation

All reads which passed the quality assessment were pooled and cross-assembled with IDBA_UD version 1.1.1 with standard parameters (Peng et al., 2012). A second dataset was added to the assembly to increase coverage (see supplementary materials & methods for details on this dataset). Prodigal v2.5 was used for prediction of protein coding DNA sequences (CDS) with the option for meta samples (Hyatt et al., 2010). Proteins were annotated with InterProScan 5.4-47.0 (Hunter et al., 2012) on the Dutch Science Grid. The annotation was further enhanced by adding EC numbers via PRIAM version March 06, 2013 (Claudel-Renard et al., 2003). Carbohydrate active modules were predicted with dbCAN release 3.0 (Yin et al., 2012). Further EC numbers were derived by text mining and matching all InterproScan derived domain names against the BRENDA database (download 13.06.13) (Chang et al., 2015). Further details on the text mining can be found in the supplementary materials & methods.

Reads were mapped back to the assembled metatranscriptome with Bowtie2 v2.0.6 (Langmead and Salzberg, 2012) using default settings. The resulting BAM files were converted with SAMtools v0.1.18 (Li et al., 2009), and gene coverage was calculated with subread version 1.4.6 (Liao et al., 2013). Read mappings to the contigs were inspected with Tablet (Milne et al., 2013). The log files with the used commands for mapping and counting can be found in supplementary file 1 and the used python script in supplementary file 2. The whole read table including all annotations can be found in supplementary file 3.

### Taxonomic assignments

All assembled contigs were analysed by blastn (Altschul et al., 1990) against the NCBI NT database (download 22.01.2014) with standard parameters, except for an e-value of 0.0001, and against the human microbiome (download 08.05.2014), the NCBI bacterial draft genomes (download 23.01.2014), the NCBI protozoa genomes (download 08.05.2014), the human genome (download 30.12.2013, release 08.08.2013, NCBI *Homo sapiens* annotation release 105) and the genomes of *Bos mutus, Bos taurus* and *Bubalus bubalis* (download 21.05.2014). Taxonomy was estimated with the LCA algorithm as implemented in MEGAN (Huson et al., 2011), but with changed default parameters. Only hits exceeding a bitscore of 50 were considered, and of these only hits with a length of more than 100 nucleotides and that did not deviate more than 10% in length from the longest hit.

For contigs, which did not retain any hits after the filtering described above, another run with blastp of the associated proteins was performed against a custom download of the KEGG Orthology (KO) database (download 25.04.2014). Taxonomic assignment was again performed with the LCA algorithm, but only hits were considered, which did not deviate by more than 10% from the hit with the maximal identity.

All taxa, which were attributed to the phylum Chordata, kingdom Viridiplantae or to artificial constructs were considered to be contaminations and were automatically removed, as well as any proteins in which the annotation contained the word “microvirus”. Furthermore, contigs that had a length of less than 300 nucleotides and which did not contain any proteins with a functional domain (disregarding the coils database) were discarded. Contigs belonging to the Illumina spike in PhiX phage were manually removed.

A compact schematic representation of the workflow is provided in Figure 1.

**Figure 1:**
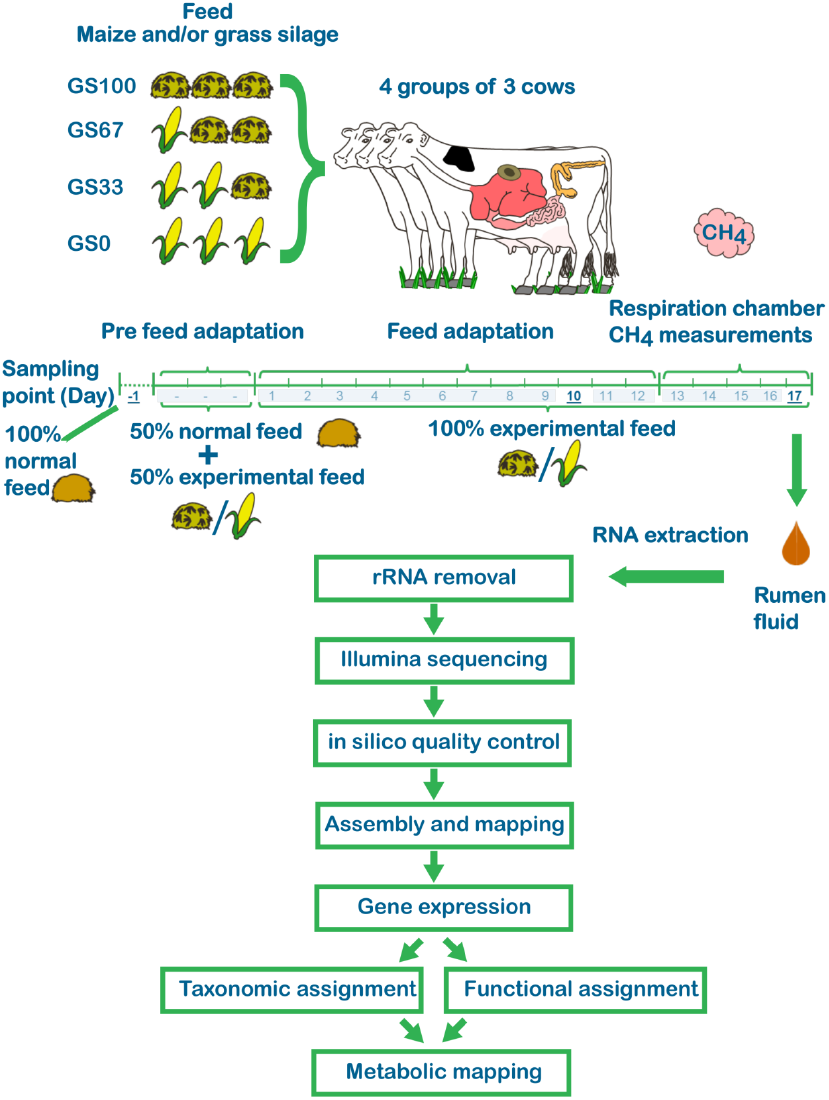
Study design. Four groups of three cows were allowed to adapt to one of four different experimental diets for twelve days. From day 13 – 17 methane emission was measured using a respiration chamber. Rumen fluid was collected at day 17 and used for microbial RNA extraction. See Methods section for details.

### Statistical analysis

Differential expression was calculated in R version 3.1.1 (R Core Team, 2012) with the edgeR package release 3.0 (Robinson et al., 2010). Only genes, which had at least 50 reads mapped in all ten samples together were considered, and only genes with a p-value and q-value <0.05 in any of the comparisons were considered to be significantly differentially expressed. Furthermore, samples from cow #14 and #511 were excluded from the statistical analysis, due to dermal antibiotic treatment and due to feeding aberrations. To examine missing links within pathways, a q-value <0.1 was also considered (referred to as “lenient approach”). The used input file, the R script with the commands, output tables and MA plots can be found in supplementary file 4. To determine whether transcription levels corresponded to the diet components, the differentially expressed genes were sorted for each gene by diet group with increasing GS content, and an increasing or decreasing isotonic regression was fit on the data. An R^2^ value of ≥0.8 was considered to be indicative of an increasing or decreasing profile, respectively, and all other values were considered to indicate that gene expression followed another, irregular, profile. Regression values and assignment of profile can be found in supplementary file 3. Isotonic regressions were computed in Python with scikit-learn version 0.15.2 (Pedregoa et al., 2011). Spearman rank correlation between the samples and Mann-Whitney U-test were calculated in Python with Scipy version 1.6.1 and NumPy version 0.9.0 (van der Walt et al., 2011).

### Metabolic mapping

All derived EC numbers were mapped with custom scripts onto the KEGG database (Kanehisa et al., 2012) and visualized using Python Scipy version 1.6.1 and NumPy version 0.9.0 (van der Walt et al., 2011) together with matplotlib version 1.4.3 (Hunter, 2007). Differentially expressed genes were investigated separately for microbial groups, which showed changes over multiple genes per pathway, and changed functions were determined by manual inspection of the KEGG maps.

### Availability of data and material

All data has been deposited at the European Nucleotide Archive (ENA) under accession numbers ERS685245 - ERS685256 and ERS710560 - ERS710568

## Results

Four experimental groups of three cows each were fed a control diet that contained only GS as roughage, and three different CS-enhanced diets for twelve days (Figure 1). From day 13 – 17, methane emission was measured using a respiration chamber, showing a significant reduction of methane emission with increasing CS proportion in the diet (van Gastelen et al., 2015). This decrease accounted for approximately 10% of the cows’ methane emission. The analysis by van Gastelen et *al*. (van Gastelen et al., 2015) showed that the dry matter feed intake of the different treatment groups did not differ significantly. Therefore the reduction in methane emission was not based on the available energy, but rather on the composition of the different diets.

Rumen fluid was collected at day 17, and used for microbial RNA extraction, mRNA enrichment and RNAseq. The complete set of RNAseq reads was cross-assembled into a single metatranscriptome. To determine activity per phylogenetic group the *de novo* assembled transcripts/genes were assigned to a taxonomic rank, and relative expression levels were obtained for four groups of animals fed different diets. Gene functional assignments were subsequently used to assess potential metabolic changes as predicted from the gene expression profiles observed in animals fed the four different diets, with a focus on carbohydrate breakdown, short chain fatty acid (SCFA) production and methane metabolism.

### Sequence, assembly and annotation metrics

In total more than 160 million reads were obtained from twelve rumen fluid samples. On average, 22.5% (Standard Deviation (SD) 6.15%) of all reads obtained per sample passed all filtering steps, retaining 18.5% of the total raw reads. Of these filtering steps, the filtering for rRNA sequences had the most impact, and removed the majority of the reads with an average of more than 63% (SD 8.75%) (all details are given in Supplementary Table 1). The majority of these rRNA reads (min. 96%) were matched to sequences from eukaryotes. The assembly yielded 712,246 contigs with in total 866,052 protein coding sequences, a length of 414,768,486 bp and an N50 of 596. While the longest contig had a size of 54,845 bp, most contigs (645,026, 90.1%) were smaller than 1000 bp. A total amount of 30 million reads, on average 58% (SD 8.75%) of the reads per sample which passed quality filtering, could be mapped back to the assembly (see Supplementary Table 1; in the following, expression values will be given relative to the amount of mapped reads, referred to as “overall expression”).

For 556,705 of the predicted protein encoding sequences a domain (excluding “Coils” domains) could be predicted. To 85,404 protein encoding sequences an EC number could be assigned.

A taxonomic classification could be obtained for 635,892 protein encoding sequences (73%), of which 282,074 could be classified at genus level. In total 1152 genera were detected, and additional 190 taxonomic assignments above the genus level were retrieved. 24 groups (at different taxonomic ranks) accounted for more than 58% of the total expression data (Figure 2). These groups included 13 genera (*Bacteroides, Butyrivibrio, Clostridium, Entamoeba, Entodinium, Eubacterium, Faecalibacterium, Fibrobacter, Methanobrevibacter, Methanosphaera, Plasmodium, Prevotella, Ruminococcus*) and 11 sequence clusters (not including the data assigned to the 13 genera) that could only be assigned at higher taxonomic levels (Archaea, Bacteria, Bacteroidales, Bacteroidetes, Clostridia, Clostridiales, Coriobacteriaceae, Eukaryota, Firmicutes, Methanobacteriaceae, Peptostreptococcaceae).

**Figure 2:**
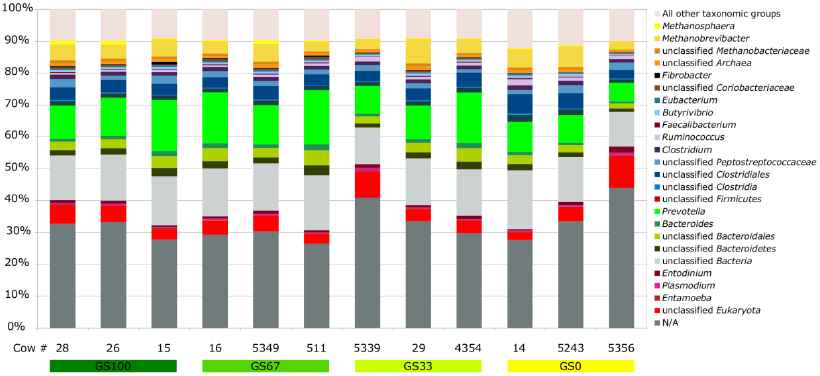
Taxonomic composition of metatranscriptome landscapes observed in animals fed one of four different diets. Diets and cows are indicated on the X-axis, taxonomic groups (at genus level, or otherwise deepest classification) are colour coded, see legend for details. N/A: No taxonomic rank could be assigned.

Fungal genes could be detected, but accounted for less than 0.1% of the overall expression. 184,991 genes without a taxonomic assignment accounted for 32% of the total expression. To only 34,731 of these genes (18.7%) any type of domain (excluding “Coils” domain) could be assigned, and only 2685 of these had an EC number assigned. Most present domains within the proteins encoded by taxonomically not assigned genes were generic domains (e.g. membrane lipoprotein attachment site, MORN repeat, P-loop containing nucleoside triphosphate hydrolase, WD40 repeat, etc.) without more specific functions.

Methanogens were represented by sequence assemblies that could be assigned to *Methanobrevibacter smithii, Methanobrevibacter ruminantium* and *Methanosphaera stadtmanae.* Reads mapping to protein coding genes assigned to methanogens captured on average 6.2% of the overall expression. In general, the overall taxonomic expression profile of the methanogens did not seem to change considerably between the different diets (Figure 2). When expression was summarized at genus level (or otherwise deepest taxonomically assigned group, as given in Figure 2, with minor groups treated together as “all other taxonomic groups”), the lowest correlation between all samples was 0.85. All microbial groups included in Figure 2 were furthermore tested (after exclusion of cows 511 and 14, due to mentioned aberrations) for statistically significant differences between animals fed the different diets (Mann--Whitney *U* test, p<0.05, not multi-test corrected), which was rejected for 150 out of 156 tests. None of the differences were statistically significant after multi-test correction (Bonferroni).

### Differential expression analysis of the rumen microbiomes

In total, 27,731 genes, which passed a set threshold for having captured at least 50 reads over all conditions combined, were subjected to the differential expression analysis, and 6397 were differentially expressed in at least one comparison (q<0.05). Three corn silage (CS) enhanced diet-induced expression profiles were distinguished (via regression analysis with isotonic regression), i.e. genes with an unlinked expression profile (profile A, 1241 genes), an induced expression corresponding to the amount of CS in the diet (profile B, 1994 genes), and a reduced expression corresponding to the amount of CS in the diet (profile C, 3162 genes) (Figure 3). Three heatmaps of all genes (per profile) can be found in supplementary file 5, displaying the overall trends within the data.

**Figure 3:**
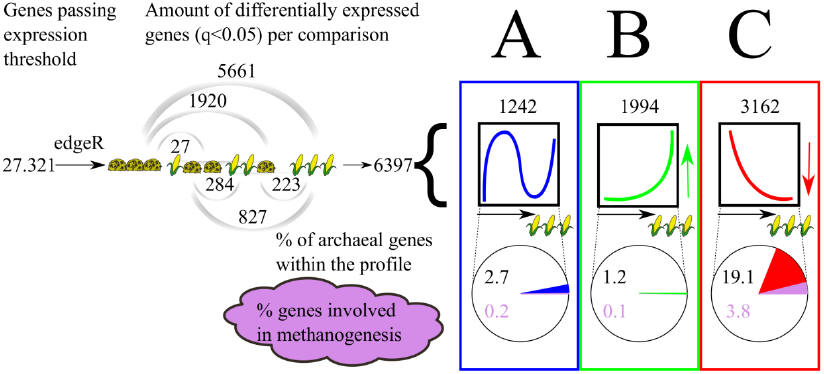
Differential expression analysis of the rumen microbiomes. Overview of the number of genes that were found to be differently expressed in pairwise comparisons of metatranscriptomes derived from animals fed different diets. Three profiles are distinguished: **Profile A**, genes with an expression, which does not follow a dietary pattern. **Profile B**, genes which are upregulated with increasing amounts of corn silage (CS). **Profile C**, genes, which are downregulated with increasing amounts of CS. Furthermore the results show that with the increase of CS, archaeal genes were mainly downregulated considerably affecting methane metabolism.

### Taxonomic and functional analysis of the three diet induced expression profiles

Genes grouped into the three different expression profiles were investigated for their taxonomic and functional classification.

For profile A, i.e. genes that did not follow a diet specific expression profile, most genes were related to general energy metabolism/carbohydrate breakdown and ribosomal protein production, as well as transport reactions. No other major functions seemed to be affected in the diet-unspecific way characteristic of profile A, and most of the genes within this group could be linked to the Clostridiales, but also to Bacteroidales, Actinobacteria and Archaea. Most predominantly represented taxa among genes following transcription profile B were bacteria belonging to the order Clostridiales, and to a lesser extent the genera Prevotella, Proteobacteria and Actinobacteria, but more than half of the differentially expressed genes could not be classified below kingdom level. The most affected functions were ribosomal protein production (mainly Eukaryota), and nucleotide metabolism in different groups, including the Eukaryota. The almost complete lack of genes associated with *Archaea* and/or methanogenesis among the genes with expression profile B indicated that there was hardly any increase of methanogenic activity with the increase of CS.

Among the genes exhibiting a lower expression upon increasing the amount of CS in the diet (profile C), the main represented microbial groups included three different methanogens (*Methanobrevibacter smithii, Methanobrevibacter ruminantium, Methanosphaera stadtmanae*), members of the genus *Prevotella*, and many genes, which could not be classified beyond the order Clostridiales. Functional profiling showed that the most downregulated processes were related to methanogenesis, electron transport and regulatory processes in the *Archaea*, as well as general metabolic functions like glycolysis, ATP generation or ribosomal protein production in all affected groups. Increased expression could also be observed for nine genes encoding putative non-ribosomal peptide synthase (NRPS) modules, among which three were taxonomically linked to *M. ruminantium* whereas the other six NRPS modules could not be classified beyond the kingdom bacteria.

With an increase of CS in the diet, Eukaryota appeared to show a decrease in their expression of genes encoding glycosylhydrolases (GH) and glycosyltransferases (GT). Furthermore, they also showed differential regulation of genes associated with movement abilities and cilia/cytoskeleton assembly, chaperons and ribosomal proteins in response to the diet changes. Most of the sequences (71.9%) assigned to the Eukaryota could not be classified below the kingdom level. For example, of the 85 differentially expressed genes encoding proteins involved in cilia/cytoskeleton assembly, only 12 could be assigned to a rank more specific than the kingdom level. Within all the classified eukaryotic sequences that showed consistent downregulation with increasing CS in the diet, the phylum Apicomplexa was the most represented, whereas the family of Ophyoscolecidae (*Entodinium, Epidinium*) showed a specific downregulation of GH encoding genes.

### Microbial starch and cellulose metabolism in cows fed with different diets

The expression of genes related to the breakdown of different complex carbohydrates differed considerably between animals fed different diets. Profile A did not include major changes in genes coding for carbohydrate degradation associated enzymes.

For genes following expression profile B, an increase of CS in the diet mainly lead to the increased expression of genes encoding different extracellular binding proteins in the genera *Ruminococcus, Bifidobacterium* and *Entodinium*, as well as an increase in the expression of genes coding for starch binding modules (CAZy classes CBM25 and CBM26) and alpha-amylases (GH13).

Most carbohydrate-metabolism associated genes affected by an increase in CS in the diet, however, followed expression profile C. With an increase of the CS in the feed, a downregulation of multiple genes involved in the breakdown of plant cell walls and their constituents could be observed, such as all the steps involved in cellulose degradation (Lynd et al., 2002). Expression of genes encoding endocellulases (CAZy classes GH5, GH9, GH45; mainly assigned to *Fibrobacter*), catalysing the first step of cellulose breakdown, was most affected, followed by genes that code for exocellulases (GH48, *Ruminococcus*) and beta-glucosidases (GH3), catalysing the second and the last step of cellulose breakdown, respectively, as well as genes encoding cellulose binding modules (e.g. CBM4, CBM13). Downregulation of the expression of genes encoding proteins involved in the breakdown of hemicellulose constituents (xylan, mannan, galactan/pectate, rhamnose) could also be observed, including genes encoding endo-1,4-beta-xylanases (GH10, GH11), beta-mannanase (GH26), pectate lyase (PL3), alpha-L-rhamnosidase (GH78), beta-1,4-galactan binding (CBM61), and xylan binding modules (e.g. CBM35). Expression of genes related to transport of glucose into the cells was also downregulated (monosaccharide transporters, EC 3.6.3.17). An overview of differentially expressed genes encoding glycosylhydrolases and carbohydrate-binding modules, including their taxonomic distribution, is presented in Figure 4 and Figure 5, respectively.

With the increase of CS, a downregulation (profile C) could be observed for *susC* and *susD* genes coding for starch binding proteins, and which could be assigned to the phylum Bacteroidetes, mainly in the genus *Prevotella*. A downregulation of expression of genes encoding proteins involved in cellulose binding was also found, including e.g. sortases, cohesins, dockerins, extracellular binding and calcium binding domains, which potentially could belong to a cellulosome (Bayer et al., 2008;Flint et al., 2008). This was mainly observed for genes assigned to the families Cellulomonadaceae, Clostridiaceae, Lachnospiraceae and Ruminococcaceae. Many functionally similar downregulated protein-coding genes could not be assigned to a taxonomic rank below the superkingdom level, mainly in the bacteria. The downregulation of a gene encoding a cohesin module was also detected in the *Archaea*, as well as the upregulation in the expression of a cohesin and dockerin module with an increase of CS.

**Figure 4:**
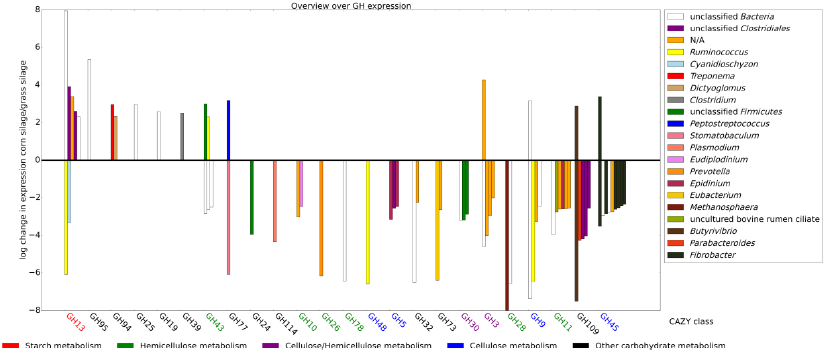
Log10 fold changes in expression of differentially expressed glycosylhydrolase encoding genes in a comparison of the 100% corn silage diet (GS0) versus the 100% grass silage diet (GS100). Positive values indicate an upregulation of gene expression in the corn silage diet. N/A: No taxonomic rank could be assigned. Colour-coding of bars indicate different taxonomic groups, whereas colour-coding of protein families indicate their involvement in the metabolism of different carbohydrates.

**Figure 5:**
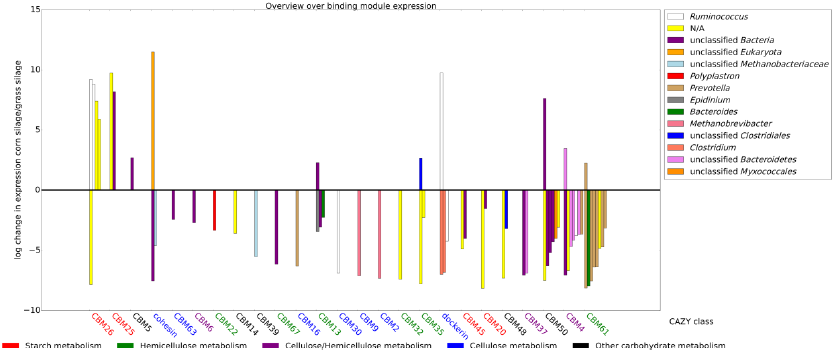
Log10 fold changes in expression of differentially expressed carbohydrate binding module encoding genes in a comparison of the 100% corn silage diet (GS0) versus the 100% grass silage diet (GS100). Positive values indicate upregulation of gene expression in the corn silage diet. N/A: No taxonomic rank could be assigned. Colour-coding of bars indicate different taxonomic groups, whereas colour-coding of protein families indicate their involvement in the metabolism of different carbohydrates.

### Microbial short chain fatty acid metabolism in cows fed different diets

The production of SCFAs is an important function of the rumen microbiome. These metabolites are taken up by the host and serve as an energy source (Bergman, 1990), have a considerable effect on methane production (van Kessel and Russel, 1996), and affect the pH, which in turn has an influence on the animal’s wellbeing (Kleen et al., 2003). In the study by Gastelen et *al*., only a small significant reduction in the SCFA butyrate was reported, with other SCFAs not changing significantly.

Increased expression upon an increase of CS in the diet (profile B) was found for genes coding for proteins which are involved in the conversion of acetyl-CoA to crotonyl-CoA, which is part of butyrate synthesis. This increase was found within the family Lachnospiraceae. The total expression of this family was on average 1.9% in all samples.

Reduced expression upon an increase of CS in the diet (profile C) was observed for genes encoding proteins involved in butyrate metabolism, and again mainly for genes assigned to the Lachnospiraceae. Several of the downregulated genes encode proteins catalysing the reactions from pyruvate to crotonyl-CoA, via acetyl-CoA, acetoacetyl-CoA and (S)-3-hydroxybutanoyl-CoA. Genes that code for enzymes catalysing the last steps to butyrate via crotonyl-CoA and butanoyl-CoA were also present in the assembly, but were not found to be differentially expressed in any of the conditions. Thus, the here presented data provide an inconclusive picture regarding the regulation of genes encoding proteins involved in ruminal butyrate production. Furthermore, consistent differential expression patterns could also not be observed for genes involved in the formation of the other SCFAs acetate or propionate. Genes encoding SCFA transporters were present in the assembly, but were not differentially expressed. Overall, these observations are in line with the fact that total SCFA concentration was found to be not affected by increasing CS in the diet, with only a minor, albeit significant increase in the molar proportion of butyrate (van Gastelen et al., 2015).

### Expression of archaeal genes involved in methane metabolism

A considerable amount of differentially expressed genes in the *Archaea* was found to encode proteins involved in methane metabolism. Based on the RNAseq data almost the complete pathways leading to methanogenesis could be reconstructed (Fig. 6). Closer inspection revealed that with an increase of CS in the diet, nearly all genes of the methanogenesis pathways were downregulated in a subset of the *Archaea* (expression profile C). Of the four possible methanogenic pathways, those for the production of methane from methanol/hydrogen, as well as from formate/carbon dioxide and hydrogen were affected. Proteins for the utilization of trimethylamines into methane could be detected in the dataset, but were not differentially expressed between animals fed the different diets. The pathway for methanogenesis from acetate was absent in the dataset.

**Figure 6:**
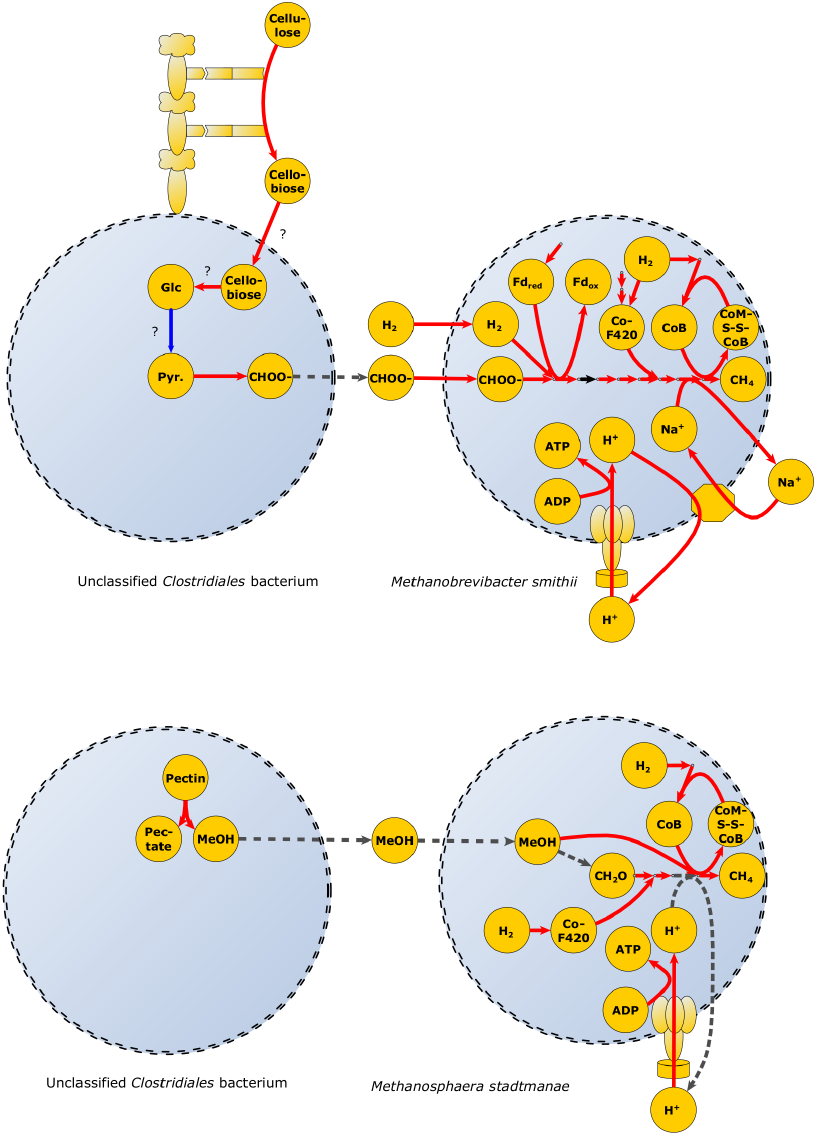
Graphical summary of metabolic consequences of the different diets in the two major methanogens and possible syntrophic partners. Red arrows: genes downregulated with the increase of corn silage in the diet; black arrows: Gene is detected but not differentially expressed; The blue arrow represents glycolysis of which the majority could be detected; punctuated arrow: orphan reactions; ? = Phylogenetic association unclear. Pyr. = Pyruvate, CHOO^-^ = Formate, FD_ox_ = oxidized Ferredoxin, FD_red_ = reduced Ferredoxin, CH_2_O = Formaldehyde.

Among genes assigned to *Methanosphaera stadtmanae,* genes coding for proteins involved in the conversion of methanol to methyl-CoM (methanol-corrinoid protein Co-methyltransferase, EC 2.1.1.90) and of methyl-CoM to methane (methyl-CoM reductase, EC 2.8.4.1) showed consistent downregulation with increasing CS in the diet (Figure 6).

Compared to changes observed for *Methanosphaera stadtmanae*, the change in the transcription pattern of genes encoding proteins involved in methanogenesis from hydrogen and formate in *Methanobrevibacter smithii* was more extensive. More specifically, the expression of genes associated with the methanogenesis pathway with formate/hydrogen was downregulated in nearly all steps (besides formylmethanofuran-tetrahydromethanopterin N-formyltransferase, EC 2.3.1.101), following expression profile C. In addition, expression of several genes encoding proteins involved in the biosynthesis of coenzyme F420 was downregulated with an increasing amount of CS in the diet (profile C). Some of these reactions could only be assigned to taxonomic levels above species, but the placements of these functions in the metabolic network indicate that they most probably can also be assigned to *M. smithii.*

Expression of genes encoding transporters for formate uptake were also downregulated (profile C), as well as genes involved in other processes related to methanogenesis, e.g. the general production of ATP, electron transport via the membrane, and sodium transport.

Nearly none of the genes that could be assigned to the third detected major methanogen in the dataset, *Methanobrevibacter ruminantium*, showed considerable downregulation, however, it should be noted that several archaeal genes, including several genes encoding proteins involved in methanogenesis, could not be classified at the species level and therefore it cannot be excluded that some of these in fact also belong to this species. Differential regulation of genes assigned to a potential syntrophic partner of *M. ruminantium, Butyrivibrio proteoclasticus* (Leahy et al., 2010), could only be detected in a few genes. Genes assigned to other formate producing organisms were also present in the data, pointing towards their potential involvement as syntrophic partners, however, no differential expression was observed for these genes, making deduction of possible syntrophic connections difficult. Further analysis of the data at the functional level showed downregulation of the expression of genes encoding proteins linked to the production of necessary substrates for methanogenesis. Expression of one of the genes encoding a subunit of pyruvate formate lyase (EC 2.3.1.54) that catalyses the production of formate from pyruvate was downregulated in a bacterium in the order Clostridiales, which could not further be classified, as well as in *Eubacterium hallii*. At the same time, several genes encoding proteins involved in the degradation of cellulose were found downregulated in animals fed CS-containing diets (profile C), and could be assigned to a not further classifiable bacterium in the order Clostridiales as well as *Ruminococcus flavefaciens, Fibrobacter succinogenes*, and several other bacteria/eukaryotes. A downregulation of genes that code for proteins involved in the production of the other substrates needed for methanogenesis, hydrogen and carbon dioxide, could not be detected.

Interestingly, using a more lenient approach (see Methods) a downregulation of expression of a gene for the production of the second major substrate for methanogenesis, methanol, was observed. More specifically, an unspecified Clostridiales bacterium showed decreased expression of a gene encoding pectinesterase (EC 3.1.1.11), catalysing the degradation of pectin to pectate and methanol.

An overview of the metabolic consequences of the observed changes in gene expression profiles is provided in Figure 6. A version of this figure with more details and a table with all reactions and assigned genes can be found in supplementary figure 1 and supplementary file 6.

## Discussion

### How feed affects methanogenesis

The rumen microbiome is a complex ecosystem, and its dynamics are determined by many variables. Most investigations to date have been focussed on the community composition and changes therein in response to different perturbations. In a recent metagenomic study by Roehe *et al*. (Roehe et al., 2016) on animals fed similar diets as the ones tested here the authors found no considerable effect on the composition of the microbiome. Here we show that in response to a diet change, gene expression within a microbiome and consequently the metabolic profile may change. Differential expression analysis revealed that although there were no extensive changes visible within the overall community expression, in line with what has previously been noticed for the sheep rumen (Shi et al., 2014), major effects could be seen regarding the expression of genes related to methane metabolism, which are also in agreement with genes which were prior identified within the metagenomics dataset by Roehe et *al*. and related publications (Wallace et al., 2015;Roehe et al., 2016;Wang et al., 2017). In two of the three methanogens identified in the dataset a coordinated downregulation of genes involved in methanogenesis as response to increased CS in the diet could be observed. Thus not only isolated single nodes involved in methanogenesis, but whole pathways were downregulated. We further found evidence for a possible syntrophy between these methanogens and several yet unidentified members of the rumen community belonging to the order of Clostridiales, which might contribute to the production of the necessary substrates (formate, methanol) for the methanogens, which was also discussed (albeit with potentially different syntrophy partners) in a related setup by Parmar et *al*. (Parmar et al., 2017). Additionally we observed a downregulation of cellulose degradation functions with increased CS in the diet. For *M. ruminatium*, we did not see a significant response to the diet changes nor did we see a significant response in possible syntrophic partners. Thus it may be that in addition to diet changes other types of biological effectors are necessary to further influence the process. Our findings are also in contrast to those reported by Shi *et al*.(Shi et al., 2014), who concluded that in the sheep rumen the supply of hydrogen is the determining factor for methane output, whereas in the present study the supply of other substrates seem to have a bigger influence. We further observed community wide responses to the change in the main energy/carbon source, with a shift in the involved glycosylhydrolases over multiple organisms and phylogenetic branches. Nevertheless, we did not observe a response in all members of the microbial community. While there was a definite downregulation of certain processes like methanogenesis, these processes were not affected in all organisms. To this end, it should be noted that the total gene count assigned to members of the Archaea greatly exceeded the size of currently known individual archaeal genomes, suggesting the presence of multiple strains of the same species in this environment (Hudman and Gregg, 1989;Sasson et al., 2017). Not all of these strains seemed to be affected by the different diets, as there were also instances of pathways, which did not show a differential regulation at all. As already observed here for the different species of methanogens, which were potentially affected because their syntrophic partners were affected, this could also be the case for the different strains of the same species, which might inhabit different niches in the rumen. It cannot be expected that e.g. methanogens living intracellularly within protozoa (Finlay et al., 1994) are in the same way affected as free living methanogens are, and that populations living closer to the substrates, i.e. those associated with the fibre fraction, will show the same behaviour as populations in the liquid fraction of the rumen (Mullins et al., 2013). Finally, as overall a reduction of methane production by ~10% was observed in this study when comparing animals fed either the GS or CS diets, it is perceivable that not all pathways and microorganisms are affected to an extend that would be detectable in significant differences in gene expression levels, also considering the relatively small sample size of three animals per experimental group.

### Unexpected findings and limitations

Several findings in this study were surprising, at least at first glance. As shown in Figure 2, and also shown by the statistical testing, the overall expression profile did not change significantly. A major change in the supplied feed was expected to result in significant changes though. Also the study of Roehe *et al*. (Roehe et al., 2016) showed no considerable changes in the relative abundance of organisms in a similar setting. We showed that the main changes are not within a taxonomic group, but rather the expression patterns per taxonomic group, which also explains the findings by (Roehe et al., 2016).

There are also concerns that differential expression analysis in communities could not reflect actual differential expression, but rather a change in organism abundance, leading to wrongly perceived changes in expression. Since in this dataset the overall expression profile per group did not statistically significantly change (although the small sampling size gives only limited power to detect this change), this is likely not an issue, and genes detected as differentially expressed are probably truly differentially expressed.

The overall taxonomic composition itself as shown in Figure 2 in general agrees with previous findings, as most of the major taxonomic groups were reported previously (Jami et al., 2013). This is also the case for the methanogens, which are similar to the ones commonly found the rumen of cows (Carberry et al., 2014) and other ruminants (Li et al., 2014;Seedorf et al., 2015). Despite this, it should be noted that the genes assigned to *Methanobrevibacter smithii* most likely belong to a related species/group of *Methanobrevibacter*, since *M. smithii* itself is not a dominant member of the rumen microbiota, but the closest sequenced relative of the species appearing in the rumen (Janssen and Kirs, 2008).

As shown in Figure 3, we also recovered changing expression profiles, which did not correspond with the diets. We were not able to find any specific functional background for these profiles, and suspect that some organisms are influenced more by the surrounding community members and not primarily by the diet, or maybe inhabit very specific niches. This would be in agreement with the findings in Figure 4 and 5, which show that a minor amount of carbohydrate active enzymes and binding modules show expression profiles against the expected trend, e.g. increase in expression of some cellulose degrading enzymes while less cellulose is fed (Özcan and Johnston, 1999;Sloothaak et al., 2015). It could also be possible that this change in expression reflects a change in metabolic strategy. As response to e.g. the lower abundance of cellulose in the environment, the affected organisms could attempt to downregulate the expression of genes coding for cellulose binding modules with low affinity, and upregulate the expression for genes coding for modules with high affinity. This mechanism is similar to the regulation of carbohydrate transporters in different organisms (Özcan and Johnston, 1999;Sloothaak et al., 2015). Additionally it needs to be considered that initial annotations might not always be correct. We found an increase in cohesin and dockerin coding modules with an increase of starch in the diet. These components are primary known as cellulosome components, but non-cellulosomal origin of these modules has been reported before (Peer et al., 2009;Ze et al., 2015). Furthermore one of these modules was found in the Archaea, which are not known to harbour either cellulosomal complexes or their starch counterparts. The same issue holds for the downregulation in expression of the genes coding for different starch binding proteins, *susC* and *susD*, which have been found to not only be starch binding, but also cellulose binding (Mackenzie et al., 2012).

Another finding, which was obvious in the investigated data, is the substantial decrease in expression of genes coding for proteins involved in cytoskeleton assembly in different Eukaryota. As several Archaea are endosymbionts of Protozoa, it can be speculated that an experimental change, which has an impact on the symbionts, will also affect their host (Finlay et al., 1994) (although this relationship is also not entirely clear (Morgavi et al., 2012)). General cellular processes, like replication, in which the cytoskeleton is involved, will then probably be directly affected, and this has been observed before in a different setting with intracellular Archaea (Holmes et al., 2014). Recently the high abundance of these proteins in the rumen proteome also have been demonstrated (Snelling and Wallace, 2017).

At last, the biggest limitation on this study are the lack of sequencing depth and little replication. The former was mainly caused by the inefficiency of the ribosomal rRNA depletion. The method used could not remove all rRNA, due to the diversity of unknown eukaryotic sequences, which resulted in a lower sequencing depth than expected. Also due to the low number of replicates, an arbitrary cutoff for the tested genes had to be applied, which is common practice and can help in some settings to increase power (Bourgon et al., 2010), and therefore it was not possible to find more subtle changes in the expression levels (e.g. a change in transcription levels of the butyrogenic pathway). Therefore this work mainly focused on changes within more highly expressed genes, and most changes were also not dependent on single p-values, but supported by expression changes in multiple genes. This has still lead to the ability to track the impact of diet on methane production, which was the aim of this study, and other effects, which were not initially expected, could still be observed. It still needs to be pointed out that the amount of replication was very small and probably too small for this type of experiment, and that many changes, including not only subtle ones, were potentially missed due to this setup.

In summary, in this study we found a significant effect of a dietary change on the gene expression in the cow rumen. A substantial fraction of the affected genes was related to methane emission, showing that a decrease in cellulose in the diet decreased the gene expression of methane related pathways. The here presented metatranscriptomic analysis is in agreement with the experimental measurements, which showed a decrease in methane emissions with the diet change (van Gastelen et al., 2015), suggesting that a change in the feed regime can have a positive effect on microbial GHG emissions.

## Acknowledgements

The authors thank Jasper Koehorst for his help with the transcriptome annotation, and Jesse van Dam, Maria Suarez-Diez and Edoardo Saccenti for helpful discussions. This computational work was carried out on the Dutch national e-infrastructure with the support of SURF Foundation.

## Author Contributions

BVDB, CMP and HS conceived designed the study. BVDB performed the experiments. BH and MD performed the bioinformatics processing and analysed the data. BH, BVDB, MD, VAPMDS, CMP, PJS and HS interpreted the results. BH, BVDB and PJS wrote the manuscript with input from the other authors. All authors read and approved the final manuscript.

### Funding

B. Hornung was supported by Wageningen University and the Wageningen Institute for Environment and Climate Research (WIMEK) through the IP/OP program Systems Biology (project KB-17-003.02-023).

This work was furthermore supported by funding from the Top Institute Food and Nutrition, Wageningen, The Netherlands, a public-private partnership on pre-competitive research in food and nutrition. The funders had no role in study design, data collection and analysis, decision to publish, or preparation of the manuscript.

## Supplementary information

### File #1

- File name: Supplementary_material_and_methods_1.docx
- File format including the correct file extension: Word document, .docx
- Title of data: Supplementary Materials and Methods
- Description of data: Extended description of the used text mining approach; processing description of the second dataset.

### File #2

- File name: Supplementary_table_1.docx
- File format including the correct file extension: Word document, .docx
- Title of data: Supplementary Table 1: Overview of the processed RNAseq data.
- Description of data: Metrics of the quality of the RNAseq data, e.g. trimming metrics

### File #3

- File name: supplementary_file_1_logs.7z
- File format including the correct file extension: 7zip archive, .7z
- Title of data: Supplementary file 1: Logfiles.
- Description of data: Logfiles of RNAseq quality control, read mapping and read counting

### File #4

- File name: supplementary_file_2_python_scripts.7z
- File format including the correct file extension: 7zip file, .7z
- Title of data: Supplementary file 2: Python scripts
- Description of data: Python scripts used for quality control, read mapping, read counting and deriving of custom EC numbers

### File #5

- File name: supplementary_file_3_all_read_counts.with_annotation.7z
- File format including the correct file extension: 7zip file, .7z
- Title of data: Supplementary file 3: Read count table with annotation
- Description of data: Read counts for the ten relevant samples, including taxonomy, functional annotation (interpro domains, EC numbers, dbcan identifiers), note if differentially expressed and regression values

### File #6

- File name: supplementary_file_4_edgeR.7z
- File format including the correct file extension: 7zip file, .7z
- Title of data: Supplementary file 4: edgeR analysis
- Description of data: input file for the edgeR analysis, R script with used commands, output tables and MA plots.

### File #7

- File name: supplementary_file_5_heatmaps.7z
- File format including the correct file extension: 7zip file, .7z
- Title of data: Supplementary file 5: Heatmaps
- Description of data: Heatmaps of all genes in the three distinguished expression patterns

### File #8

- File name: supplementary_file_6.methane_related_processes_fig_6_curated.7z
- File format including the correct file extension: 7z file, .7z
- Title of data: Supplementary file 6: Overview over methane related reactions.
- Description of data: Overview over methane related reactions with assigned genes, corresponding to figure 6/supplementary figure 1.

### File #9

- File name: supplementary_figure_1_rumen_metabolism_with_legend.svg
- File format including the correct file extension: scalable vector graphic, .svg
- Title of data: Supplementary figure 1: Annotated version of figure 6.
- Description of data: Annotated version of figure 6, with details given in supplementary file 6.

